# Klp2 and Ase1 synergize to maintain meiotic spindle stability during metaphase I

**DOI:** 10.1101/2020.01.31.929729

**Authors:** Fan Zheng, Fenfen Dong, Shuo Yu, Tianpeng Li, Yanze Jian, Lingyun Nie, Chuanhai Fu

## Abstract

The spindle apparatus segregates bi-oriented sister chromatids during mitosis but mono-oriented homologous chromosomes during meiosis I. It has remained unclear if similar molecular mechanisms operate to regulate spindle dynamics during mitosis and meiosis I. Here, we employed live-cell microscopy to compare the spindle dynamics of mitosis and meiosis I in fission yeast cells and demonstrated that the conserved kinesin-14 motor Klp2 plays a specific role in maintaining metaphase spindle length during meiosis I, but not during mitosis. Moreover, the maintenance of metaphase spindle stability during meiosis I requires the synergism between Klp2 and the conserved microtubule crosslinker Ase1 as the absence of both proteins causes exacerbated defects in metaphase spindle stability. The synergism is not necessary for regulating mitotic spindle dynamics. Hence, our work reveals a new molecular mechanism underlying meiotic spindle dynamics and provides insights into understanding differential regulation of meiotic and mitotic events.

## INTRODUCTION

During meiosis, duplicated chromosomes are segregated into daughter cells to form haploid gametes through two rounds of successive cell division (Ohkura, 2015). The spindle apparatus is responsible for chromosome segregation, and both centrosome-dependent and centrosome-independent mechanisms have been reported to regulate spindle assembly (Ohkura, 2015; Petry, 2016). Despite the diverse mechanisms used to regulate spindle assembly, the spindle dynamics of meiosis and mitosis is remarkably similar. Particularly, both meiotic and mitotic cells require their spindle lengths to be constant during metaphase in order to promote chromosome alignment for further segregation (Goshima & Scholey, 2010). Motor and/or microtubule crosslinking proteins have been found to function antagonistically within the spindle to maintain metaphase spindle length (Blackwell et al., 2017; Civelekoglu-Scholey, Tao, Brust-Mascher, Wollman, & Scholey, 2010; Goshima & Scholey, 2010; Pidoux, LeDizet, & Cande, 1996; Rincon et al., 2017; W. Saunders, Lengyel, & Hoyt, 1997; W. S. Saunders & Hoyt, 1992; Syrovatkina, Fu, & Tran, 2013; van Heesbeen, Tanenbaum, & Medema, 2014; Yukawa, Yamada, Yamauchi, & Toda, 2018). Due to the lack of comparative studies of meiotic and mitotic spindles, it has remained unclear whether the metaphase spindle length in meiotic and mitotic cells is regulated by similar mechanisms.

It is conceivable that specific molecular mechanisms may operate during meiosis I to regulate spindle dynamics because the spindle mechanics of meiotic and mitotic cells is different. During mitotic metaphase, kinetochores on sister chromatids are bi-oriented and captured by microtubules from the opposite spindle poles, whereas, during metaphase of meiosis I, kinetochores on a homologous chromosome are mono-oriented and captured by microtubules from the same spindle pole (Ohkura, 2015). This special configuration of kinetochores during meiosis I requires chiasmata, instead of cohesion, to provide physical links between two homologous chromosomes to resist the separating forces contributed by meiotic spindles.

The evolutionarily conserved kinesin-5 and kinesin-14 motor proteins have been shown to play important roles in regulating spindle assembly and dynamics. Generally, kinesin-5 functions to promote spindle bipolarization during preanaphase and works with kinesin-14 in an antagonistic manner to maintain the bipolar spindle during metaphase (Civelekoglu-Scholey et al., 2010; Pidoux et al., 1996; Rincon et al., 2017; W. Saunders et al., 1997; W. S. Saunders & Hoyt, 1992; She & Yang, 2017; Yukawa et al., 2018). Additionally, kinesin-14 works in concert with the conserved Ase1/PRC1/MAP65 family proteins to crosslink microtubules and to organize antiparallel microtubule arrays (Braun et al., 2011; Carazo-Salas & Nurse, 2006; Daga, Lee, Bratman, Salas-Pino, & Chang, 2006; Janson et al., 2007). Since discovered in *Drosophila* (Endow, Henikoff, & Soler-Niedziela, 1990), kinesin-14/NCD has been shown to play critical roles in spindle pole focusing, spindle assembly and stabilization, and chromosome segregation during meiosis and mitosis (Endow, Chandra, Komma, Yamamoto, & Salmon, 1994; Hallen, Liang, & Endow, 2008; Hatsumi & Endow, 1992a, 1992b; Komma, Horne, & Endow, 1991; Matthies, McDonald, Goldstein, & Theurkauf, 1996; Skold, Komma, & Endow, 2005; Theurkauf & Hawley, 1992). Similarly, the kinesin-14 motors in *Xenopus* (XCTK2) (Walczak, Verma, & Mitchison, 1997) and human (HSET) (Cai, Weaver, Ems-McClung, & Walczak, 2009; Mountain et al., 1999) play important roles in proper spindle assembly. Extensive biochemical characterizations establish that kinesin-14 is a slow possessive motor possessing a microtubule crosslinking activity (Braun, Drummond, Cross, & McAinsh, 2009; Braun et al., 2017; Fink et al., 2009; Furuta & Toyoshima, 2008; Janson et al., 2007; Walker, Salmon, & Endow, 1990) (Reinemann, Norris, Ohi, & Lang, 2018). Interestingly, a NCD mutation was reported to affect meiotic, but not mitotic, chromosome segregation (Komma et al., 1991). Moreover, inhibition of HSET activity compromises meiotic spindle organization in marine oocytes but does not appear to affect mitotic spindle architecture in cultured cells (Mountain et al., 1999). These findings not only underscore the evolutionarily conserved roles of kinesin-14 in organizing microtubule arrays and regulating spindle assembly but also highlight different roles of kinesin-14 in meiotic and mitotic cells. The different role of kinesin-14 in meiosis and mitosis remains to be further investigated.

The fission yeast *Schizosaccharomyces pombe* is an excellent model organism for studying meiosis due to the ease of obtaining meiotic cells and the convenience of monitoring spindle and chromosome dynamics by live-cell microscopy (Fu et al., 2009; Yamashita, Sakuno, Watanabe, & Yamamoto, 2017). Two kinesin-14 motors, i.e. Pkl1 and Klp2 (Troxell et al., 2001), are present in fission yeast. Pkl1 localizes mainly at the spindle pole body to anchor microtubule minus ends (Syrovatkina & Tran, 2015; Yukawa, Ikebe, & Toda, 2015), whereas Klp2 functions to organize antiparallel microtubule arrays at microtubule plus ends during interphase and to regulate kinetochore dynamics at kinetochores during mitosis (Carazo-Salas & Nurse, 2006; Gachet et al., 2008; Grishchuk & McIntosh, 2006; Janson et al., 2007). Genetically, both Pkl1 and Klp2 oppose the function of Cut7/kinesin-5 (Troxell et al., 2001), and spindle bipolarity restores if Pkl1 is removed in cells lacking Cut7 (Olmsted, Colliver, Riehlman, & Paluh, 2014; Syrovatkina & Tran, 2015). The restoration of spindle bipolarity in cells lacking both Pkl1 and Cut7 depends on the microtubule bundling protein Ase1 and microtubule associated proteins Alp7 and Alp14 (Rincon et al., 2017; Yukawa et al., 2017). Unlike Pkl1 and its counterpart in *Drosophila* (i.e. NCD), the role of Klp2 in mitotic spindle assembly is ill-defined and the role of klp2 in regulating meiotic spindle dynamics has not been determined.

We employed live-cell microscopy to study spindle dynamics of mitosis and meiosis I in a comparative manner and demonstrated that the conserved kinesin-14 motor Klp2 plays a crucial role in preventing spindle collapse during meiosis I, but not during mitosis. Similar to Ase1, Klp2 may function as a microtubule crosslinker and synergizes with Ase1 to maintain spindle stability during meiosis I. Our work therefore provides important insights into understanding the differential regulation of meiotic and mitotic spindle dynamics.

## RESULTS

### The spindles of meiosis I and mitosis display characteristic dynamics

During meiosis I, chiasmata form between homologous chromosomes and the kinetochores on sister chromatids are mono-oriented. These characteristics of meiosis promoted us to hypothesize that the spindle of meiosis I may be regulated specially in order to properly segregate the homologous chromosomes. To test this hypothesis, we first employed live-cell microscopy to analyze the spindle dynamics of mitosis and meiosis I in fission yeast cells expressing Sid4-GFP (a protein marking spindle pole body) and mCherry-Atb2 (α-tubulin). As reported previously (Nabeshima et al., 1998; Yamamoto et al., 2008), the spindle dynamics of mitosis and meiosis I exhibits three characteristic phases: phase I (prophase-like, during which the spindle assembles), phase II (metaphase/anaphase A, during which the spindle length remains relatively constant), and phase III (anaphase B, during which the spindle length increases rapidly) (Figure 1, A-C). Despite the similarity, the two types of spindle dynamics displayed different characteristics. As shown in Figure 1D, the spindle elongated faster during phase I of meiosis I (referred to as prophase I) than during phase I of mitosis and maintained a longer length during phase II of meiosis I (referred to as metaphase I) than during phase II of mitosis. By contrast, the duration of phase I and II of meiosis I (referred to as preanaphase I) and mitosis was comparable (Figure 1D).

**Figure 1.**
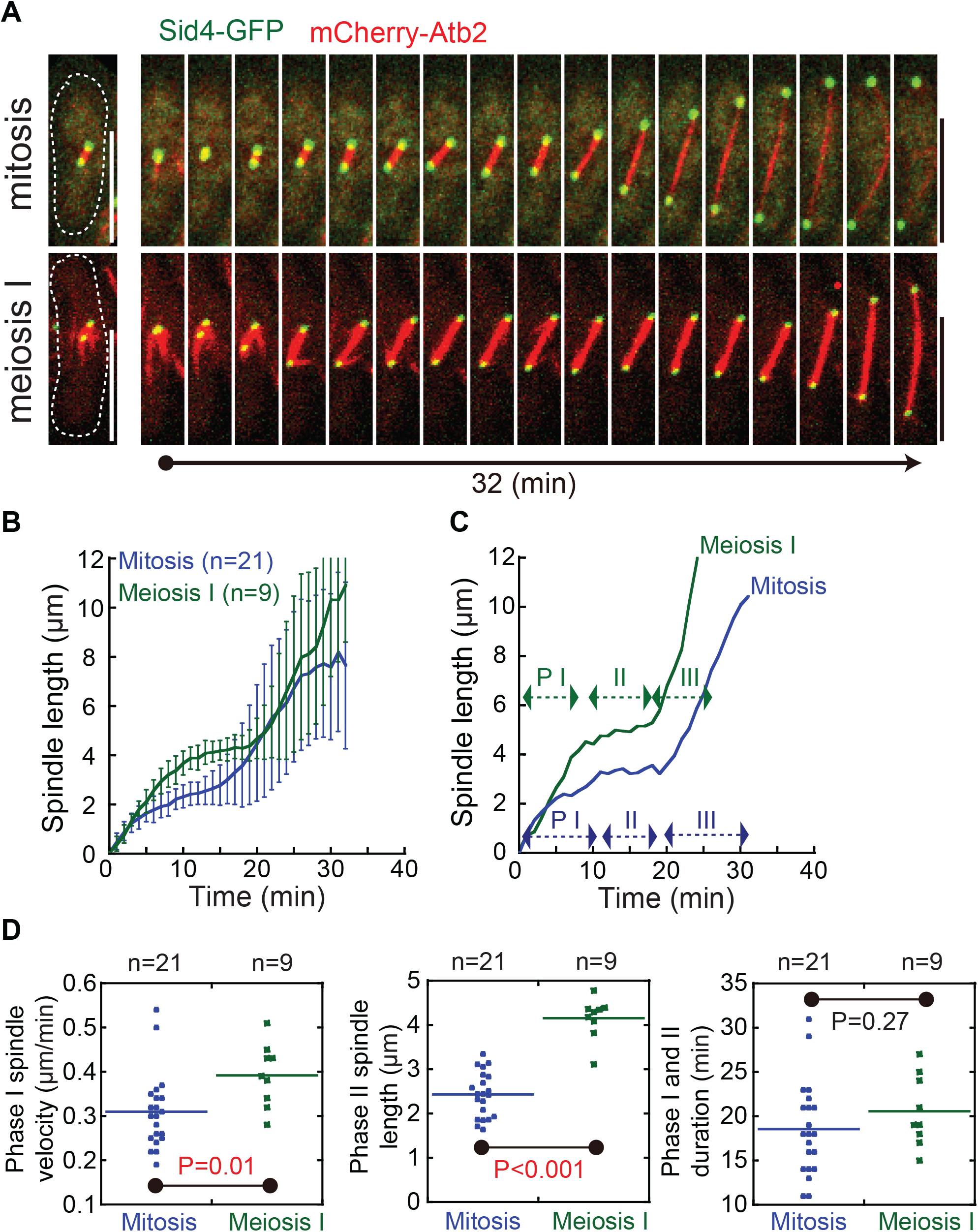
Spindle dynamics during meiosis I and mitosis in wild type cells. (A) Maximum projection time-lapse images of wild type (WT) cells expressing Sid4-GFP (a protein marking the spindle pole body) and mCherry-Atb2 (α-tubulin) in mitosis and meiosis I. Dashed lines mark cell outline. Scale bar, 10 μm. (B) Plots of spindle length against time for the cells in mitosis and meiosis I. The initial time point is the time when spindle microtubules emerged. Error bars represent standard deviation, and cell number analyzed is indicated. (C) Representative plots of spindle dynamics for the cells in mitosis and meiosis I. Phase I (prophase-like), phase II (metaphase), and phase III (anaphase B) are indicated. (D) Quantification of prophase spindle speed, metaphase spindle length, and preanaphase duration for mitotic and meiotic cells. *p* values were calculated by students’ t-test, and cell number analyzed is indicated.

### Screening for motor proteins involved in regulating spindle dynamics of meiosis I

We then attempted to identify the proteins that are responsible for regulating meiotic spindle dynamics. We focused on motor proteins because they are one of the main types of proteins involved in spindle assembly and regulation (Reber & Hyman, 2015). Live-cell microscopy was employed to examine meiotic spindle dynamics of dynein-deletion cells (*Dhc1Δ*) and kinesin-deletion cells (all nonessential kinesins). As shown in supplementary Figure S1, most of the kinesin/dynein mutants displayed defective spindle dynamics during meiosis I, but to different extents, with *klp2Δ* (kinesin-14 in human), *klp5Δ* (kinesin-8 in human), and *klp6Δ* (kinesin-8 in human) cells showing the strongest phenotypes. In the absence of Klp5 or Klp6, most of the meiotic spindles continuously elongated and did not show a distinguishable metaphase I. Interestingly, in the absence of Klp2, about half of the meiotic spindles abruptly collapsed during metaphase I (Supplementary Figure S1 and Figure 2), suggesting that Klp2 functions mainly to maintain spindle stability during metaphase I. Therefore, in this present study, we addressed the role of Klp2 in regulating meiotic spindle dynamics.

**Figure 2.**
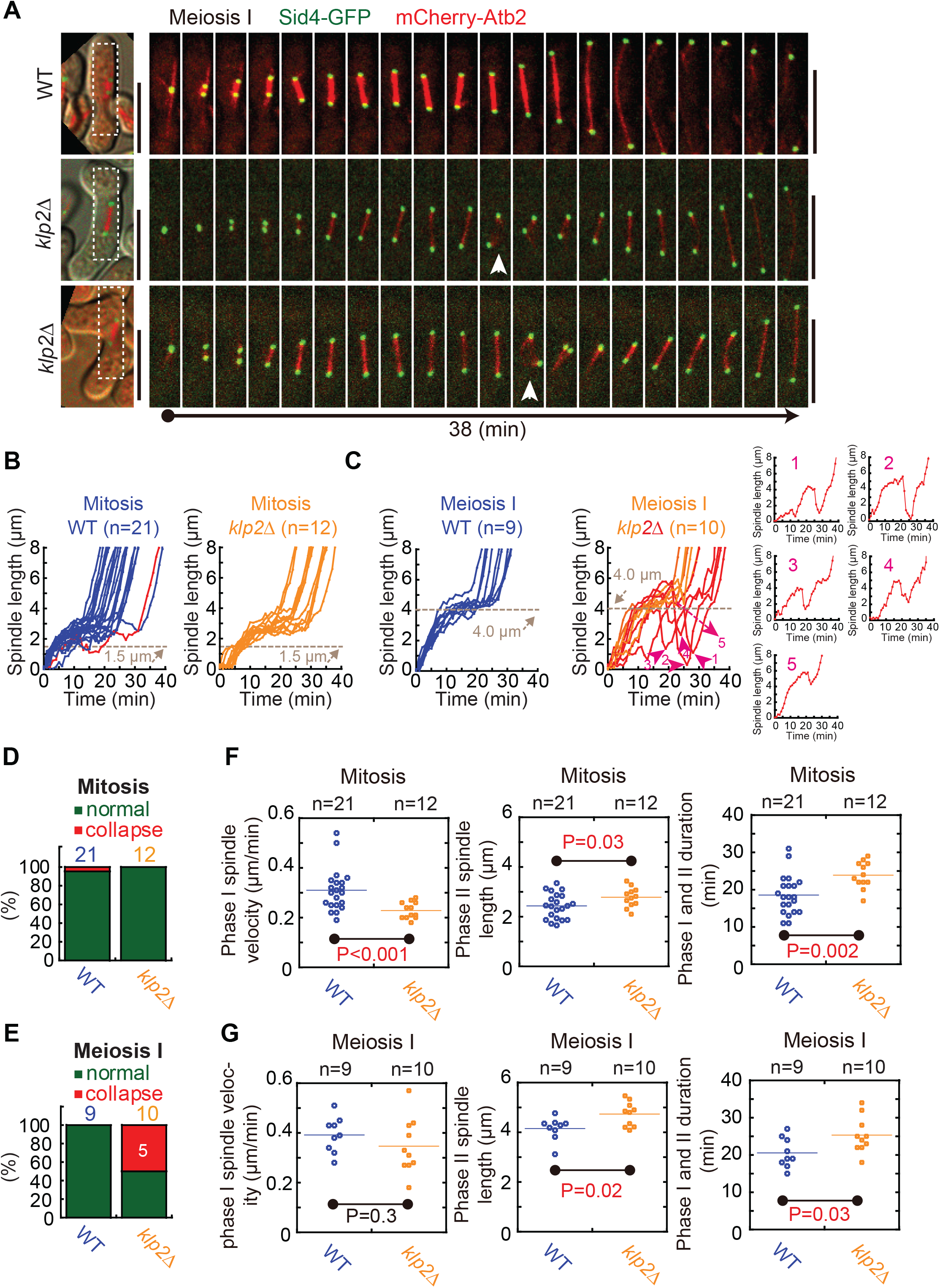
*klp2Δ* cells display spindle collapse during meiosis I, but not during mitosis. (A) Maximum projection time-lapse images of meiotic WT and *klp2Δ* cells expressing Sid4-GFP and mCherry-Atb2. Arrowheads indicate the collapsing spindles. Dashed rectangles are the selected regions used for displaying the time-lapse images. Scale bar, 10 μm. (B) Plots of spindle length against time for mitotic WT and *klp2Δ* cells. Cell number analyzed is indicated. Note that both types of cells displayed no spindle collapse except one WT cell showed slight spindle regression (highlighted in red). The data of WT cells is the same as the one used in Figure 1, B-D. (C) Plots of spindle length against time for meiotic WT and *klp2Δ* cells. Cell number analyzed is indicated. Note that half of the spindles in *klp2Δ* cells collapsed abruptly during metaphase I (numbered and shown individually). The data of WT cells is the same as the one used in Figure 1, B-D. (D and E) Quantification of spindle collapse for WT and *klp2Δ* cells in mitosis (D) and meiosis I (E). Cell number analyzed is indicated, and the white number indicates the number of the cells displaying spindle collapse. (F and G) Quantification of prophase spindle speed (phase I), metaphase spindle length (phase II), and preanaphase duration (phase I and II) for WT and *klp2Δ* cells in mitosis (F) and meiosis I (G). *p* values were calculated by students’ t-test, and *n* indicates cell number analyzed.

### Klp2 is required for preventing spindle collapse during metaphase I

For comparison, we examined spindle dynamics of WT and *klp2Δ* cells during mitosis and meiosis I. Mitotic spindle dynamics of WT and *klp2Δ* cells was comparable (Figure 2B), and no spindle collapse was detected except that one WT cell (21 in total) displayed slight spindle regression during metaphase (Figure 2D). By contrast, 50% of the spindles in *klp2Δ* cells underwent abrupt collapse during metaphase I (Figures 2A, 2C and 2E). The spindle collapse may not be due to the abnormalities of spindle elongation and length because the absence of Klp2 did not significantly affect spindle elongation during prophase I and only slightly lengthened the maximal spindle length during metaphase I (Figure 2G). Further analysis of mitotic spindle parameters showed similar results except that the absence of Klp2 slowed down spindle elongation during prophase (Figure 2F). In both mitotic and meiotic cells, the absence of Klp2 slightly but significantly prolonged the duration of preanaphase (Figures 2F and 2G). These results not only show similar functions of Klp2 in regulating spindle dynamics but also highlight the meiosis-specific role of Klp2 in preventing spindle collapse.

### Klp2 is required for maintaining proper organization of the interpolar microtubules during meiosis I

Next, we employed live-cell microscopy to image WT and *klp2Δ* cells expressing Bub1-GFP (a spindle assembly checkpoint protein localized to kinetochores) and mCherry-Atb2. As shown in Figure 3A, interpolar and kinetochore microtubules were distinguishable in the *klp2Δ* cells undergoing spindle collapse. Specifically, the absence of Klp2 leaded to buckling of the interpolar microtubules in concomitant with rapid shortening of the distance between the two spindle poles (Figure 3A). Consistent with the quantification data shown in Figure 2E, spindle collapse was detected in >50% of the Bub1-GFP-expressing *klp2Δ* meiotic cells but no spindle collapse was found in the Bub1-GFP-expressing *klp2Δ* mitotic cells (Figure 3B).

**Figure 3.**
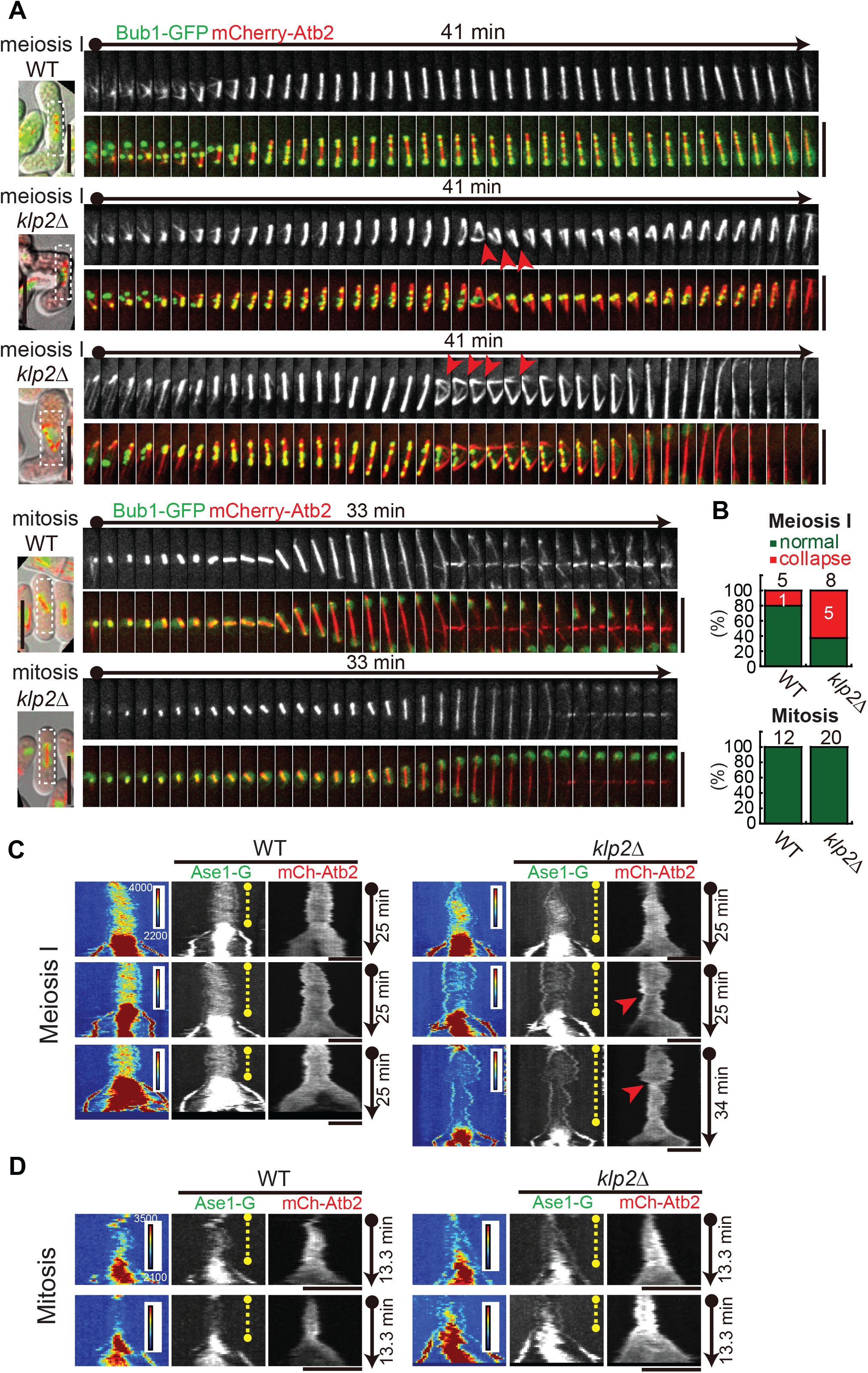
The absence of Klp2 affects proper localization of Ase1 to the spindle and causes spindle buckling in meiotic cells. (A) Maximum projection time-lapse images of WT and *klp2Δ* cells expressing Bub1-GFP and mCherry-Atb2. Red arrowheads mark the collapsing spindles in meiotic cells. Dashed rectangles are the selected regions used for displaying time-lapse images. Note that no mitotic cells displayed spindle collapse. Scale bar, 10 μm. (B) Quantification of spindle collapse for WT and *klp2Δ* cells in mitosis and meiosis I. Cell number analyzed is indicated, and the white number indicates the number of the cells displaying spindle collapse. (C and D) Kymograph analysis of Ase1 and spindle dynamics of WT and *klp2Δ* cells in meiosis I (C) and mitosis (D). Ase1 signal intensity is shown by using hitmap graphs. Yellow dashed lines indicate preanaphase while red arrowheads mark the collapsing spindles. Scale bar, 5 μm.

Since Ase1 (PRC1 in human) is a crucial factor crosslinking the antiparallel interpolar microtubules (Fu et al., 2009; Janson et al., 2007), we further assessed the effect of the absence of Klp2 on the localization of Ase1 within the spindle. Intriguingly, we noticed that Ase1 signals within the spindle during preanaphase were much brighter in meiotic cells than in mitotic cells (Figure 3C and 3D), suggesting that more Ase1 molecules are concentrated on the preanaphase spindle, presumably within the antiparallel interpolar microtubules, during meiosis I than during mitosis. Moreover, the absence of Klp2 impaired the localization of Ase1 within the spindle during preanaphase of meiosis I in the *klp2Δ* cells displaying spindle collapse, but not in the *klp2Δ* cells displaying normal spindle dynamics (Figure 3C). By contrast, the localization of Ase1 within the spindles in mitotic WT and *klp2Δ* cells appeared to be comparable (Figure 3D). Together, these results suggest that Klp2 plays a crucial role in organizing the interpolar microtubules within the meiotic spindles during preanaphase, likely through regulating the localization of Ase1.

### Overexpression of Ase1 rescues spindle collapse in klp2Δ cells during meiosis I

We reasoned that overexpression of Ase1 in cells lacking Klp2 could restore the localization of Ase1 to the interpolar microtubules and subsequently rescue spindle collapse. To test this hypothesis, we expressed GFP-Klp2 and Ase1-GFP from the *ase1* promoter in *klp2Δ* cells and performed live-cell microscopy to analyze spindle dynamics during meiosis I (Figure 4A). Note that Ase1-GFP-expressing *klp2Δ* cells contain both ectopic GFP-Ase1 and endogenous Ase1. The expression of the GFP-tagged proteins was confirmed in both h+ and h− cells (Figure 4B), which were crossed for analyzing spindle dynamics during meiosis I. As shown in Figures 4C and 4D, meiotic spindle dynamics in GFP-Klp2-expressing or Ase1-GFP-expressing *klp2Δ* cells and in WT cells (see Figure 2C) was comparable, and only 10% and 20% of GFP-Klp2-expressing and Ase1-GFP-expressing *klp2Δ* cells, respectively, displayed spindle collapse (vs. 50% in *klp2Δ* cells shown in Figure 2E). Therefore, ectopic expression of both GFP-Klp2 and Ase1-GFP largely rescued spindle collapse caused by the absence of Klp2. Hence, it is possible that Klp2 and Ase1 synergize to organize interpolar spindle microtubules and maintain spindle length during metaphase I.

**Figure 4.**
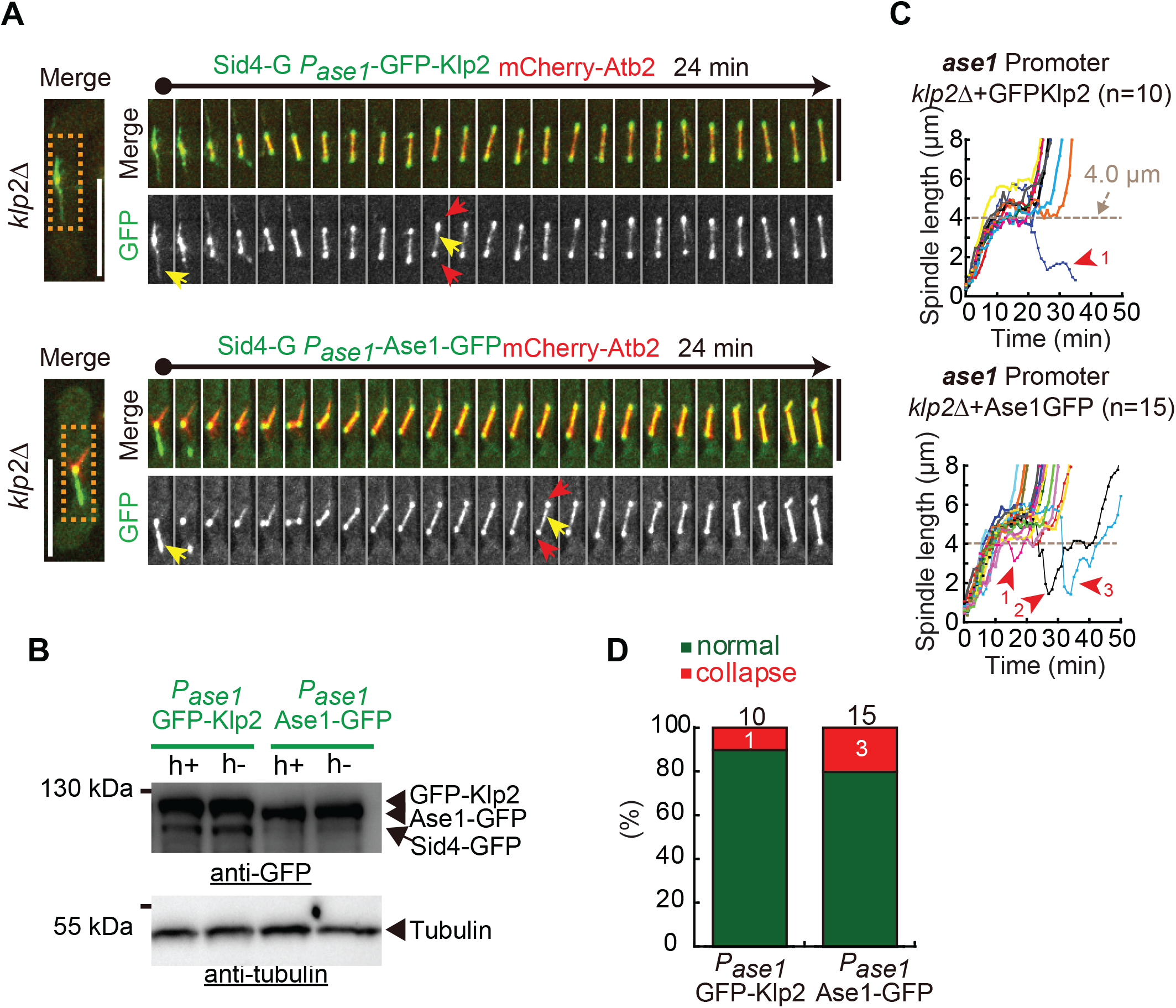
Overexpression of Ase1 rescues spindle collapse in *klp2Δ* cells. (A) Maximum projection time-lapse images of *klp2Δ* cells expressing mCherry-Atb2, Sid4-GFP, and *P_ase1_*-GFP-Klp2 (from the *ase1* promoter) or *P_ase1_*-Ase1-GFP (from the *ase1* promoter). Red arrows mark the SPBs while yellow arrows indicate the localization of GFP-Klp2 or Ase1-GFP. Dashed rectangles are the selected regions used for displaying time-lapse images. Scale bar, 5 μm. (B) Western blotting performed to test the expression of GFP-Klp2 or Ase1-GFP in h+ and h− cells with antibodies against GFP and tubulin. (C) Meiotic spindle dynamics of *klp2Δ* cells expressing *P_ase1_*-GFP-Klp2 or *P_ase1_*-Ase1-GFP. Cell number observed for analysis is indicated. Spindle collapse is marked by numbered red arrowheads and dashed lines mark a spindle length of 4.0 μm. (D) Quantification of spindle collapse in GFP-Klp2 or Ase1-GFP-expressing *klp2Δ* cells. Cell number analyzed is indicated and the white number indicates the number of the cells displaying spindle collapse.

### Ase1 is required for maintaining spindle stability during metaphase I

To reveal the role of Ase1 in regulating spindle dynamics, we analyzed spindle dynamics during mitosis and meiosis I in *ase1Δ* cells by live-cell microscopy. In mitotic *ase1Δ* cells, the maximal metaphase spindle length was significantly reduced as compared with the one in mitotic WT and *klp2Δ* cells (Figures 5A and 5B and Figure 2B). In 3 of the 17 mitotic *ase1Δ* cells, the spindles did not appear to elongate before anaphase onset. These results suggest a critical role of Ase1 in promoting spindle elongation during mitotic prophase. By contrast, in meiotic *ase1Δ* cells, the maximal metaphase spindle length was comparable with the one in meiotic WT and *klp2Δ* cells (Figures 5C and 5D and Figure 2C) but the spindles appeared to be destabilized with the length oscillating/regressing during metaphase I. Given the main role of Ase1 in organizing interpolar microtubules (Braun et al., 2011; Janson et al., 2007), it was surprising to see that the spindles in *ase1Δ* cells were able to elongate and did not collapse during meiosis I. This finding suggests that in addition to Ase1, other factors such as Klp2 may be required for maintaining spindle stability during metaphase I.

**Figure 5.**
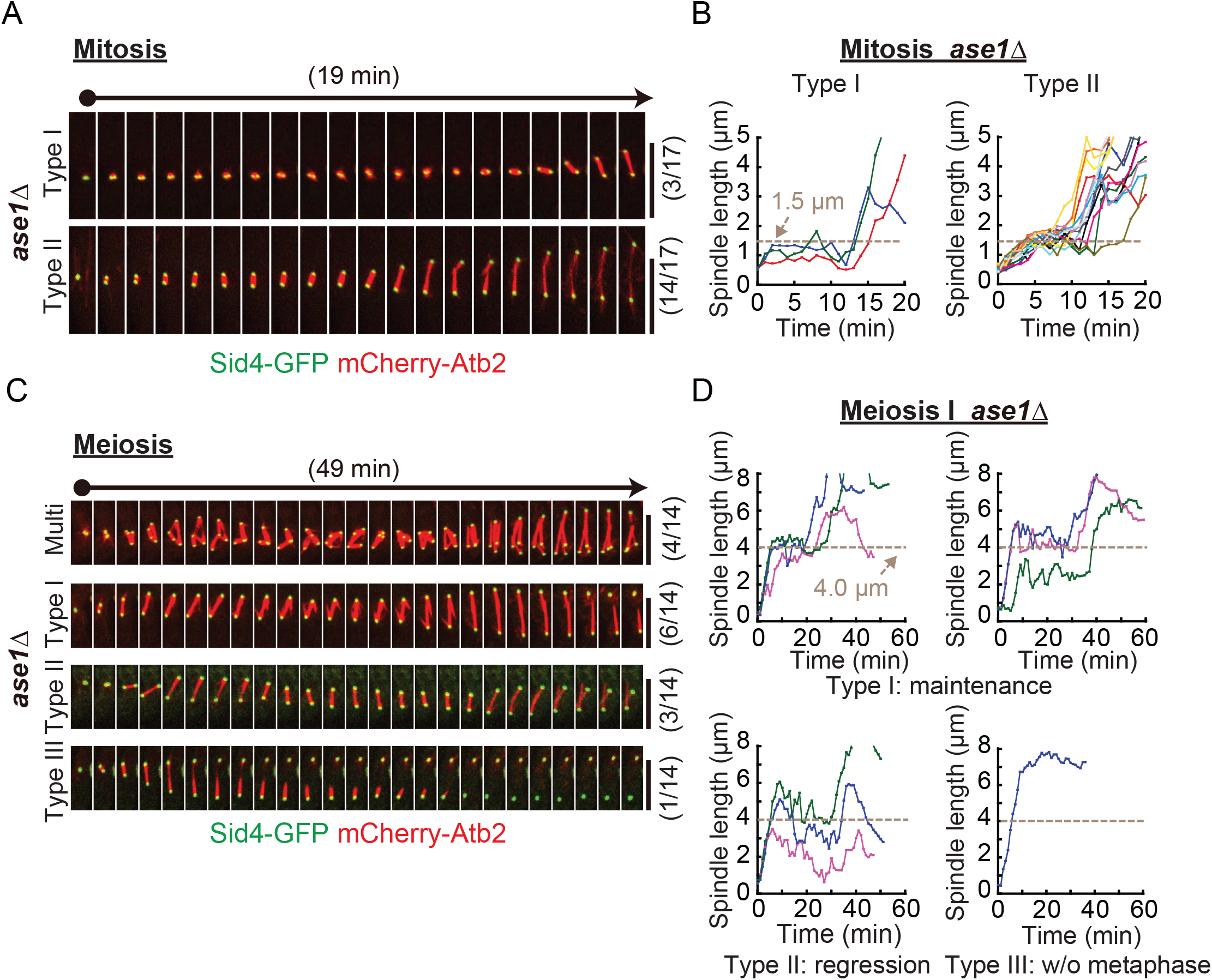
The absence of Ase1 destabilizes spindles during meiosis I and causes short metaphase spindles during mitosis. (A) Maximum projection time-lapse images of mitotic *ase1Δ* cells expressing Sid4-GFP and mCherry-Atb2. Two typical types of spindle dynamics are shown (see explanation in (B)). The number of cells observed for each type is indicated in brackets. Scale bar, 5 μm. (B) Plots of spindle length against time of mitotic *ase1Δ* cells indicated in (A). Two typical types of curves are shown: I) lacking spindle elongation during prophase and II) shortened metaphase spindle length. Note that the dashed lines indicate a spindle length of 1.5 μm. (C) Maximum projection time-lapse images of meiosis I *ase1Δ* cells expressing Sid4-GFP and mCherry-Atb2. A multipolar spindle and three typical types of spindles are shown (see explanation in (D)). The number of cells observed for each type is indicated in brackets. Scale bar, 5 μm. (D) Plots of spindle length against time of meiotic *ase1Δ* cells (indicated in (C)). Three typical types of plots are shown: I) wild-type-liked metaphase spindle length (maintenance), II) spindle regression (regression), and III) continuous spindle elongation (lacking metaphase). Note that the dashed lines indicate a spindle length of 4.0 μm.

### Klp2 and Ase1 work in concert to maintain spindle stability during metaphase I

To understand the interplay between Klp2 and Ase1 in regulating spindle dynamics, we then assessed the spindle dynamics of mitosis and meiosis I in *ase1Δklp2Δ* cells. Intriguingly, no additive spindle defect was found in *ase1Δklp2Δ* cells during mitosis and the spindles in *ase1Δklp2Δ* cells did not elongate properly and were short (Figures 6A and 6B). By contrast, an additive effect on spindle destabilization was detected in *ase1Δklp2Δ* cells during meiosis I (Figures 6C and 6D). Specifically, the spindles in *ase1Δklp2Δ* cells failed to maintain a stable metaphase length during meiosis I and spindle microtubules in most of the *ase1Δklp2Δ* cells (5/7) were found to splay apart. These results suggest a synergistic role of Ase1 and Klp2 in organizing interpolar spindle microtubules and maintaining spindle stability during metaphase I.

**Figure 6.**
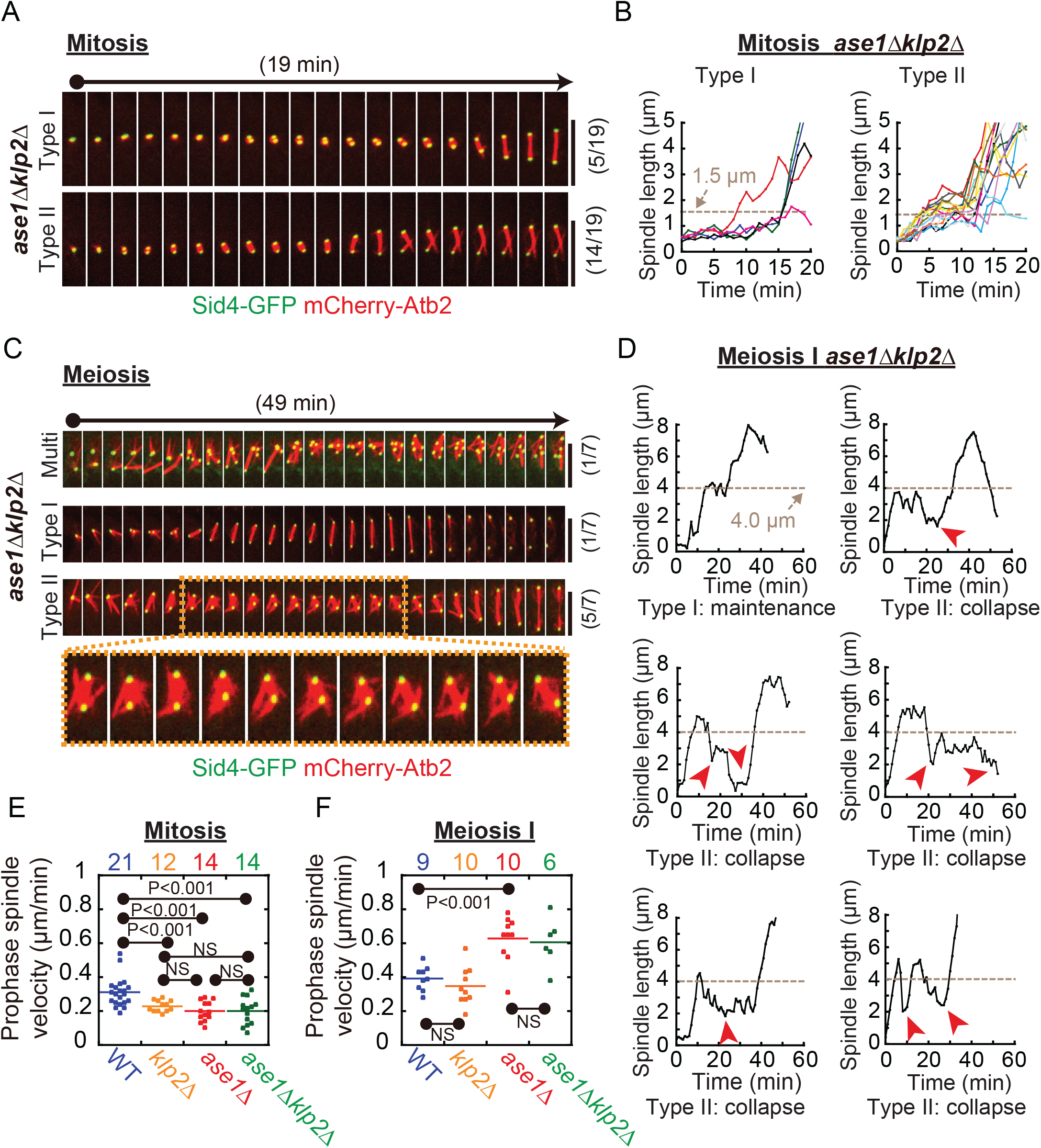
The absence of both Ase1 and Klp2 exacerbates spindle destabilization during meiosis I. (A) Maximum projection time-lapse images of mitotic *ase1Δklp2Δ* cells expressing Sid4-GFP and mCherry-Atb2. Two typical types of spindle dynamics are shown (see explanation in (B)). The number of cells observed for each type is indicated in brackets. Scale bar, 5 μm. (B) Plots of spindle length against time of mitotic *ase1Δklp2Δ* cells indicated in (A). Two typical types of curves are shown: I) lacking spindle elongation during prophase and II) shortened metaphase spindle length. Note that the dashed lines indicate a spindle length of 1.5 μm. (C) Maximum projection time-lapse images of meiotic *ase1Δklp2Δ* cells expressing Sid4-GFP and mCherry-Atb2. Of the seven cells analyzed, one displays multiple spindle poles (indicated as “Multi”) and one and five display the type-I and -II spindle phenotypes, respectively, indicated in (D). Magnified inset images highlight the splaying of spindle microtubules. Scale bar, 5 μm. (D) Plots of spindle length against time of meiotic *ase1Δklp2Δ* cells. Two typical types of plots are shown: I) wild-type-liked metaphase spindle length (maintenance), and II) spindle collapse (collapse). Note that red arrowheads mark spindle collapse. (E) Quantification of mitotic prophase spindle speed. The speed was determined by measuring the slope of the prophase curves after linear regression. Cell number analyzed is indicated, and “N.S.” means non-significant. The *p* values were calculated by students’ t-test. Note that the data for WT and *klp2Δ* mitotic cells, shown in Figure 2F, is also shown for comparison. (F) Quantification of meiotic prophase spindle speed. The speed was determined by measuring the slope of the prophase curves after linear regression. Cell number analyzed is indicated, and “N.S.” means non-significant. The *p* value was calculated by students’ t-test. Note that the data for WT and *klp2Δ* meiotic cells, shown in Figure 2G, is also shown for comparison.

It is worth pointing out that Ase1 may also play a role in promoting spindle bipolarity because a small portion of *ase1Δ* cells (4/14) (Figure 5C) and *ase1Δklp2Δ* cells (1/7) (Figure 6C), but not WT (0/9) and *klp2Δ* (0/10) cells, displayed multipolar spindles.

We further assessed the spindle elongation during mitotic and meiotic prophase in *ase1Δ* and *ase1Δklp2Δ* cells. Intriguingly, the spindle elongated more slowly during mitotic prophase in *ase1Δ*, *klp2Δ*, and *ase1Δklp2Δ* cells than in WT cells (Figure 6E). This result suggest that Ase1 and Klp2 are required for promoting spindle elongation during mitotic prophase. By contrast, the spindle elongated much faster during meiotic prophase in *ase1Δ* and *ase1Δklp2Δ* cells than in WT and *klp2Δ* cells (Figure 6F). This finding establishes that Ase1 plays a special role during prophase I in restricting spindle elongation. Collectively, Klp2 and Ase1 are required for promoting spindle elongation during mitotic prophase but synergize to maintain spindle stability during metaphase I.

### Klp2 and Ase1 are required for proper segregation of homologous chromosomes

Finally, we dissected chromosome segregation during meiosis I and mitosis. As shown in Figure 7, A-D, a range of abnormalities in homologous centromere segregation during meiosis I were found in *ase1Δ*, *klp2Δ*, *klp2Δase1Δ*, and even WT cells. Based on the degree of severity, the abnormalities were classified into two categories, in which one displayed unsegregated centromeres after spindle breakdown (i.e. centromere lagging and clustering upon spindle breakdown) and the other displayed segregated centromeres that ultimately converged at the two SPBs upon spindle breakdown (Figure 7E). Interestingly, 7.7% (n=13) of WT meiotic cells displayed unsegregated centromeres, suggesting that homologous chromosome segregation is error-prone during meiosis I (Figure 7E). The absence of Ase1 or Klp2 enhanced defects in homologous chromosome segregation with 38.1% (n=21) and 28.6% (n=14) of the *klp2Δ* and *ase1Δ* cells, respectively, displaying unsegregated centromeres (Figure 7E). Moreover, 43.7% (n=16) of the *klp2Δase1Δ* cells displayed unsegregated centromeres (Figure 7E). This quantification result correlated well with the degree of spindle defects caused by the absence of Klp2 and/or Ase1 (Figures 2C, 5D, and 6D). Thus, Klp2 and Ase1 are involved in regulating homologous chromosome segregation, likely through ensuring proper maintenance of meiotic stability.

**Figure 7.**
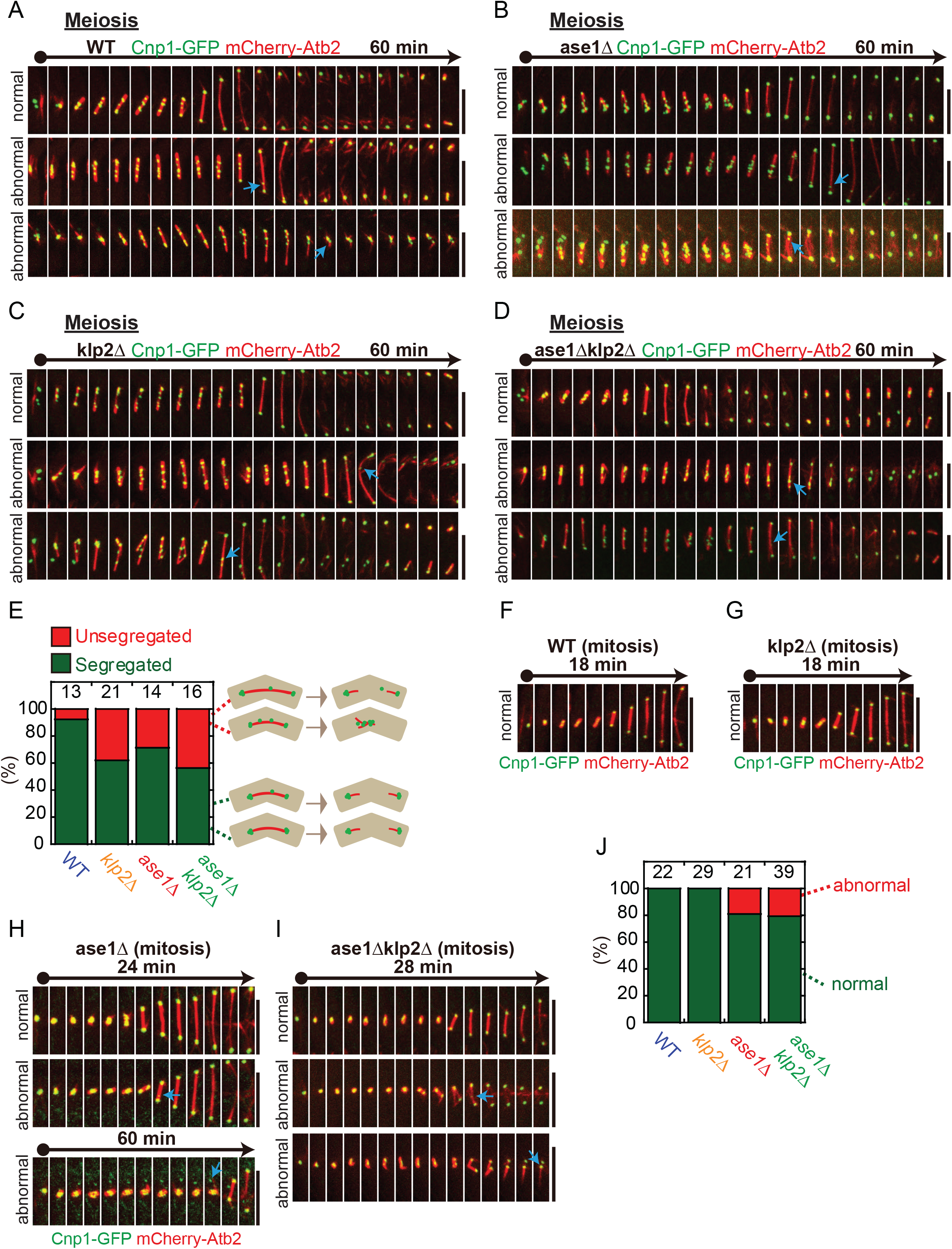
Centromere dynamics in WT, *ase1Δ*, *klp2Δ*, and *ase1Δklp2Δ* cells. (A-D) Maximum projection time-lapse images of WT (A), *ase1Δ* (B), *klp2Δ* (C), and *ase1Δklp2Δ* (D) meiotic cells expressing Cnp1-GFP and mCherry-Atb2. Arrows mark the lagging or clustering centromeres. Scale bar, 5 μm. (E) Quantification of the meiotic cells displaying unsegregated centromeres. Cell number analyzed is indicated. (F-I) Maximum projection time-lapse images of mitotic WT (F), *klp2Δ* (G), *ase1Δ* (H), and *ase1Δklp2Δ* (I) cells expressing Cnp1-GFP and mCherry-Atb2. Arrows mark abnormally segregated centromeres. Scale bar, 5 μm. (J) Quantification of abnormally segregated centromeres for the mitotic cells. Cell number analyzed is indicated.

For comparison, we also examined centromere segregation during mitosis. As shown in Figure 7, F-J, a small portion of cells lacking Ase1 displayed abnormal centromere segregation whereas the absence of Klp2 did not affect mitotic chromosome segregation.

## DISCUSSION

Few studies were focused on addressing the difference between mitotic and meiotic spindle dynamics. In this present work, we compared mitotic and meiotic spindle dynamics in a quantitative manner and showed that kinesin-14/Klp2 plays a crucial role in regulating spindle and chromosome dynamics during meiosis I, but not during mitosis. We further demonstrated that Klp2 works in concert with Ase1 to organize interpolar spindle microtubules during meiosis I and to mediate homologous chromosome segregation. Hence, our work reveals a new meiotic mechanism underlying the maintenance of metaphase spindle length.

Our comparison analysis elucidated that the spindle dynamics of mitosis and meiosis I exhibits distinct characteristics (Figure 1). The spindle length of meiotic I phase II is ~2-fold longer than that of mitosis (Figures 1B an 1D), in consistence with the findings reported previously (Nabeshima et al., 1998; Yamamoto et al., 2008). By contrast, the preanaphase duration of mitosis and meiosis I are comparable (Figure 1D). Therefore, the spindle elongation speed during phase I of meiosis I is significantly faster than that of mitosis (Figures 1B and 1D). Klp2 does not appear to contribute to the increased speed of meiotic spindle elongation (Figure 2G). Instead, Klp2 plays a specific role in maintaining metaphase spindle length during meiosis I, but not during mitosis, since about half of the spindles in *klp2Δ* cells collapse abruptly during meiosis I (Figures 2A and 2C).

The abrupt collapse of metaphase spindles in *klp2Δ* cells is likely due to the defective organization of interpolar microtubules (Figure 3A). It has been reported that the budding yeast orthologue of Klp2 (i.e. Kar3) aligns interpolar microtubules to the spindle axis so that Cin8/kinesin-5 can efficiently slide apart the antiparallel microtubules to promote spindle bipolarity during mitosis (Hepperla et al., 2014). Therefore, the absence of Kar3 causes short metaphase spindles during mitosis. However, the absence of the Kar3 counterpart Klp2 in fission yeast slightly but significantly lengthens metaphase spindle during mitosis and meiosis I (Figures 2F and 2G). Therefore, it is unlikely that Klp2 regulates meiotic spindle dynamics through enhancing the functionality of kinesin-5. We noticed that the localization of Ase1, the microtubule-crosslinker organizing the antiparallel interpolar microtubules within the spindle (Fu et al., 2009; Janson et al., 2007; Loiodice et al., 2005; Rincon et al., 2017; Yamashita, Sato, Fujita, Yamamoto, & Toda, 2005), to interpolar microtubules is impaired in *klp2Δ* cells during meiosis I (Figure 3C). This finding suggests the possibility that Klp2 regulates meiotic spindle dynamics through facilitating the localization of Ase1 to spindle microtubules. This is further supported by the fact that artificial enhancing the localization of Ase1 to meiotic spindles by Ase1 overexpression can rescue spindle collapse in *klp2Δ* cells (Figure 4).

Consistent with previous studies (Braun et al., 2011; Janson et al., 2007), the synergism between Klp2 and Ase1 is required for proper organization of meiotic spindles. This is supported by the facts that the absence of both Klp2 and Ase1 exacerbates spindle destabilization during preanaphase of meiosis I (Figures 6C and 6D) and that *klp2Δase1Δ* cells display defective spindle organization with splayed microtubules (Figure 6C). The synergism mechanism may not operate in mitotic cells to regulate metaphase spindle stability because the absence of Klp2 and Ase1 does not causes spindle collapse during mitosis (Figures 6A and 6B). It is worth pointing out that less Ase1 localizes to interpolar microtubules during preanaphase of mitosis than that of meiosis I and that the absence of Klp2 does not appear to affect the localization of Ase1 in mitotic cells (Figures 3C and 3D). Nevertheless, consistent with the previous reports (Loiodice et al., 2005; Yamashita et al., 2005), our work confirmed that Ase1 is required for promoting spindle elongation during mitotic preanaphase (Figures 5A and 5B).

Surprisingly, Ase1 is not required for promoting spindle elongation during meiotic preanaphase I. Instead, the absence of Ase1 significantly increases the preanaphase spindle speed of meiosis I (Figure 6F). Moreover, the absence of Ase1 does not significantly shorten the maximum length of metaphase spindles during meiosis I (Figures 5D). These findings demonstrate the differential roles of Ase1 in regulating mitotic and meiotic spindle dynamics. We hypothesize that Ase1 may function in a dose-dependent manner within the spindle. When the concentration of Ase1 is elevated within the spindle during meiosis I, presumably in a Klp2 dependent manner (Figure 3C), Ase1 functions as a limiting factor in restricting spindle elongation. When the levels of Ase1 are low within the spindle during mitosis, Ase1 then serves as a stabilizing factor to promote spindle elongation (Figure 3D). This hypothesized model awaits further investigation.

Why do meiotic spindles need both Ase1 and Klp2, but not Ase1 alone, to maintain metaphase spindle stability? It has been established that like Ase1, Klp2 family proteins also function as microtubule crosslinkers (Braun et al., 2017; Ludecke, Seidel, Braun, Lansky, & Diez, 2018; Peterman & Scholey, 2009). Klp2 and Ase1 have been shown to work in concert to organize antiparallel microtubule arrays (Braun et al., 2011; Janson et al., 2007). Therefore, it is conceivable that a more sophisticated mechanism involving multiple types of microtubule crosslinkers (i.e. Ase1 and Klp2) is required to construct and maintain metaphase spindles of a larger size, e.g. meiotic spindles. Meiotic spindles elongate faster than mitotic spindles during preanaphase (Figure 1D). We postulate that the involvement of Ase1 and Klp2 could also provide more robust forces, likely entropic forces as shown previously (Braun et al., 2017; Lansky et al., 2015), to antagonize spindle sliding and subsequently to maintain the stability of meiotic metaphase spindles.

In conclusion, this present work reveals a meiosis-specific role of Klp2 in maintaining metaphase spindle stability and provides insights into understanding differential regulations of meiotic and mitotic spindle dynamics.

## MATERIALS and METHODS

### Plasmids and yeast strains

Strains were created by random spore digestion or tetra dissection. Plasmids were constructed by homologous recombination using the ClonExpress®II One Step Cloning Kit (www.vazyme.com). The strains and plasmids used in this study are listed in the supplementary tables 1 and 2, respectively.

### Microscopy

Imaging was performed with a PerkinElmer UltraVIEW Vox spinning-disk microscope equipped with a Hamamatsu C9100-23B EMCCD camera and a CFI Apochromat TIRF 100x objective (NA=1.49). Cells were sandwiched between an EMM (Edinburgh Minimal Medium)-agarose pad and a coverslip for imaging, as reported previously (Shen et al., 2019). For mitosis analysis, vegetative cells were grown in EMM media plus five supplements (Adenine, Leucine, Uracil, Histidine, and Lysine, 0.225 g/L each) and were collected for imaging until OD600 reached 0.5-1.0. For meiosis I analysis, cells with opposite mating types were first streaked on YE (Yeast extract) plates and were grown overnight at 30 °C. The fresh cells with opposite mating types from the YE plates were then mixed on ME (Malt extract) plates for 10 hours before the cells were collected for imaging. All media were purchased from Formedium (www.formedium.com). Briefly, stack images containing 11 planes at 0.5 μm spacing were acquired every minute. Time-lapse maximum projection images were created using MetaMorph 7.7 (www.moleculardevices.com).

### Data analysis

Spindle length was determined by measuring the distance between two spindle poles using imaging J 1.5 (imagej.nih.gov). Graphs of spindle length against time were then created using KaleidaGraph 4.5 (www.synergy.com) to show spindle dynamics. Phase I (prophase-like) and phase III (anaphase B) are the phases showing continuous spindle elongation. During phase II (metaphase), the spindle length slightly oscillates around its mean. Phase I spindle speed (i.e. prophase spindle speed) was determined using linear regression that was applied to the spindle length plots for Phase I. Phase II spindle length (i.e. metaphase spindle length) was determined by calculating the mean spindle length of phase II. In the case of spindle collapse, phase II without the phase of collapsing was used for the calculation. The duration of phase I and II (i.e. preanaphase) was the time between the initiation of spindle assembly and anaphase B onset. All plots were created using KaleidaGraph 4.5. Kymograph graphs were constructed using MetaMorph 7.7.

## ACKNOWLEDGEMENTS

We thank Dr Phong Tran (UPENN) and the Yeast Genetic Resource Center Japan for providing yeast strains and plasmids.. This work is supported by grants from National Key Research and Development Program of China (2017YFA0503600), National Natural Science Foundation of China (31671406, 91754106, 31871350, and 31621002), the Strategic Priority Research Program of the Chinese Academy of Sciences (XDB19040101), and China’s 1000 Young Talents Recruitment Program.

## AUTHOR CONTRIBUTIONS

C.F. conceived and supervised the project. F.Z., F.D., S.Y., T.L., Y.J., and L.N. performed experiments. F.Z., S.Y., and C.F. analyzed data. C.F. wrote the paper. All authors made comments.

## Supplemental Material

### Supplemental figure

**Figure S1.**
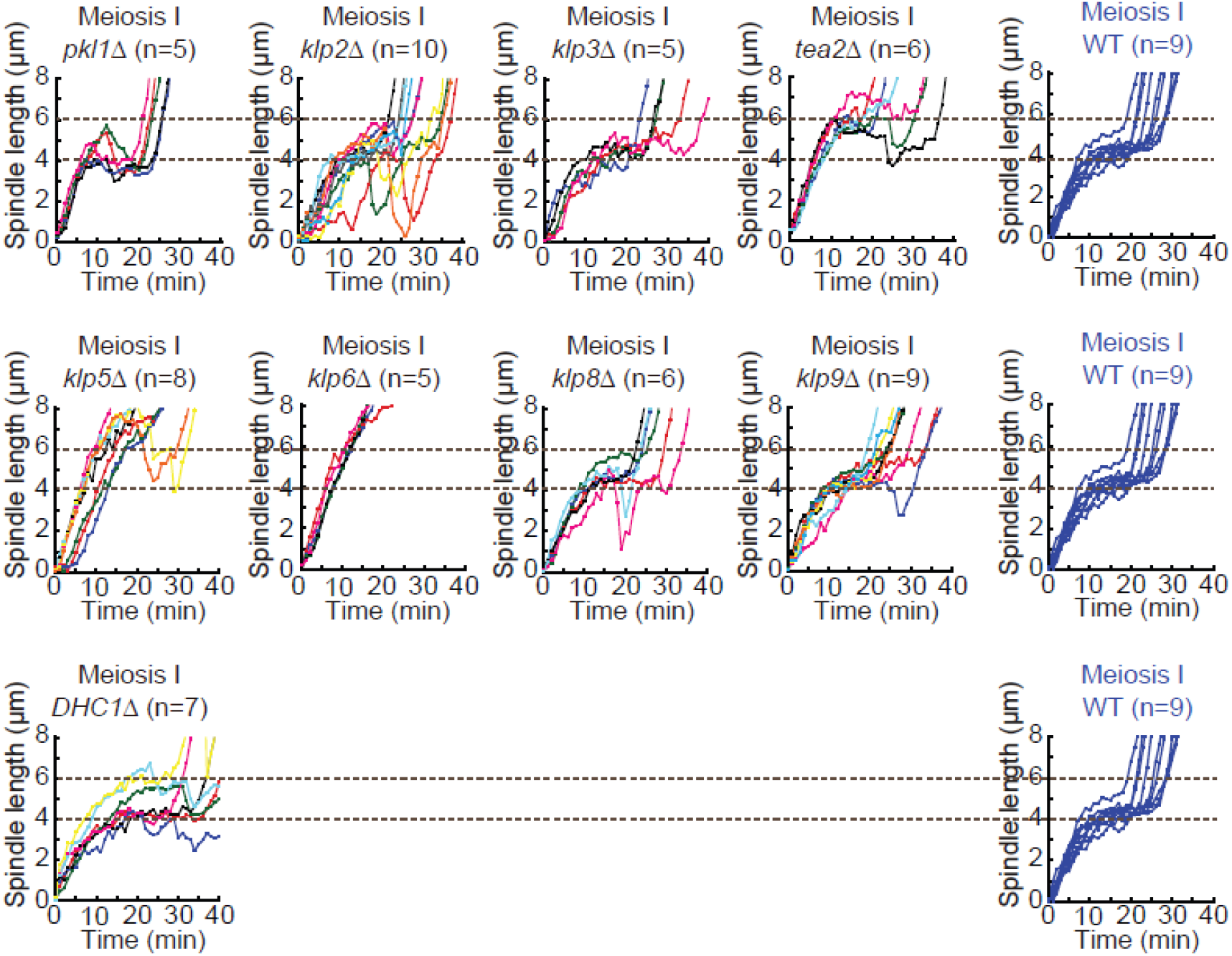
Spindle dynamics during meiosis I in the indicated cells. Cell number analyzed was indicated. Two dashed lines indicate 4 and 6 μm, respectively.

### Supplemental Tables

**Table S1.**
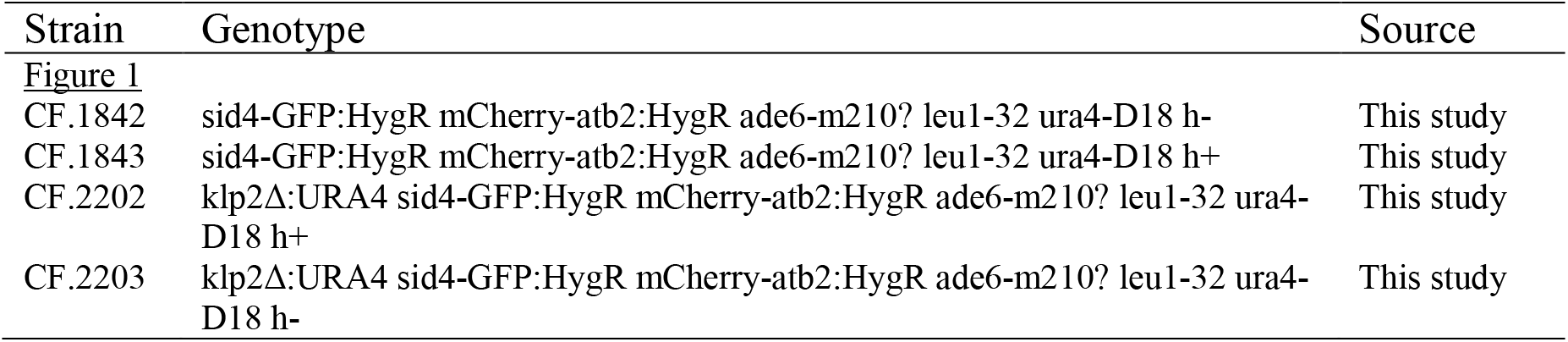
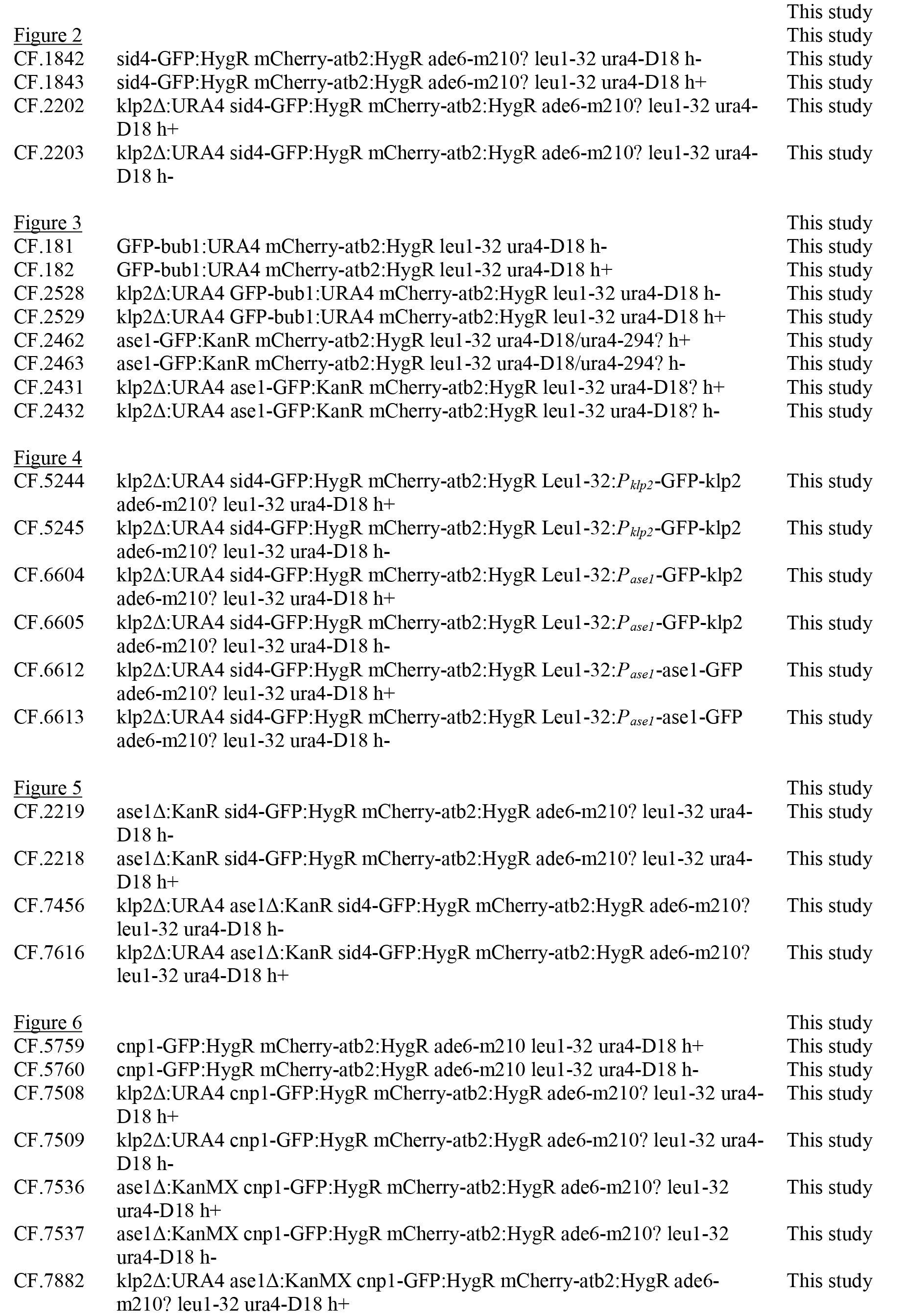
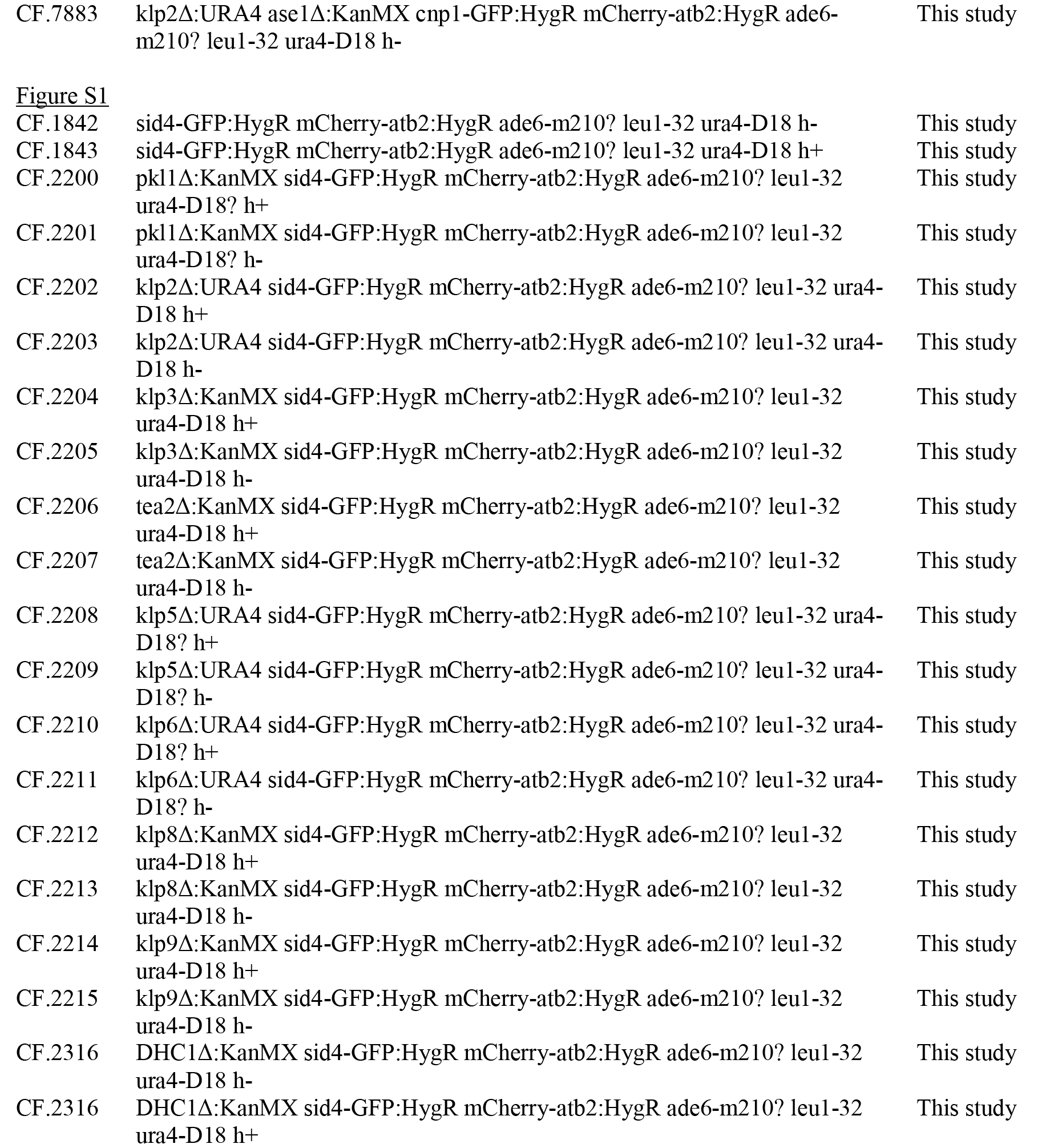
Yeast strains

**Table S2.** Plasmids

| Plasmid | Genotype | Source |
| --- | --- | --- |
| **Figure 4** |  |  |
| pCF.2062 | pJK148-*P<sub>klp2</sub>*-GFP-klp2 | This study |
| pCF.1926 | pJK148-*P<sub>ase1</sub>*-GFP-klp2 | This study |
| pCF.861 | pJK148-*P<sub>ase1</sub>*-ase1-GFP | This study |

